# Conservation genetics of regionally extinct peregrine falcons (*Falco peregrinus*) and unassisted recovery without genetic bottleneck in southern England

**DOI:** 10.1101/2020.04.08.031914

**Authors:** Angela Weaving, Hazel A. Jackson, Michael K. Nicholls, Jon Franklin, Rodrigo Vega

## Abstract

The peregrine falcon (*Falco peregrinus*) has been affected by persecution, pollution, trade and habitat degradation, but it is considered a flagship conservation success story because of successful reintroductions. However, in the UK there were never formal reintroduction programmes for peregrine falcons, and it appears that UK populations – and specifically the Sussex peregrines of the English south coast – recently recovered from a population crash unassisted. To study this, we obtained samples from contemporary populations in southern England, Ireland, continental Europe, domestic-bred peregrine falcons, and from England pre-population crash. Using microsatellite and mtDNA control region data, the genetic diversity and structure, signatures of genetic bottlenecks, and potential origin of the Sussex peregrines was investigated. We found low levels of genetic diversity across all peregrine falcon populations, low but significant genetic differentiation among all populations, and a few private alleles, indicating some level of genetic structure in European peregrines. Although we could not pinpoint the origin of the Sussex peregrines, the data suggests that it is not likely to have originated from escaped domestic birds or from adjacent European populations. The results obtained here parallel other studies on peregrines elsewhere showing low genetic diversity but genetic structure. We conclude that not enough time elapsed for genetic erosion to occur due to the population bottleneck, and that at least for the Sussex peregrines there is no need for genetic conservation by wild-take and subsequent captive breeding programmes as long as current protection measures remain in place.

## Introduction

The peregrine falcon (*Falco peregrinus*, Tunstall 1771) has a wide distribution and it can be found in all continents except Antarctica (White 1994). Although it is considered as a Least Concern species by the International Union for the Conservation of Nature (IUCN), the peregrine falcon is a charismatic species, admired for its speed, agility and hunting behaviour (Ratcliffe 2010), which has been severely affected by persecution, pollution, trade and habitat degradation (BirdLife International 2016). The peregrine falcon is now considered a flagship conservation success story because of effective reintroductions, e.g. Canada, United States of America, Southern Scandinavia, Germany and Poland aiding its recovery from localised extinctions (Crick and Ratcliffe 1995; Ratcliffe 2003; Banks et al. 2010; Smith et al. 2015). However, although in the UK there were never formal reintroduction programmes for peregrine falcons, populations in southern England recovered from a population crash unassisted (Franklin and Everitt 2009). This brings up the question as to where do the re-established peregrines in southern England come from and have they suffered the detrimental impacts associated with a genetic bottleneck?

During the middle of the 20^th^ century, peregrine falcon populations in many regions of the northern hemisphere, including mainland UK, underwent dramatic declines (Crick and Ratcliffe 1995; Horne and Fielding 2002; Ratcliffe 2010). Prior to 1939, Sussex (Southern England) supported the densest breeding population in any part of the British Isles (Ratcliffe 1963; unpublished diaries John Walpole-Bond 1904-1954). The wartime (1939-1945) persecution of peregrines, due to them being predators of homing pigeons (war or carrier pigeons) carrying military messages, saw many coastal UK peregrine populations decline (Ratcliffe 1963). The small number of fatalities in Sussex however, had no persisting effect on the local population. The number of breeding pairs recorded immediately after the war between 1945 and 1951 are comparable with the records for the years leading up to 1940. After World War II, the main reasons for the peregrine falcon population ‘crash’ included direct persecution of adults (Humphreys et al. 2007; Ratcliffe 2010) and indirect mortality of adults and eggshell thinning due to persistent organochlorine pesticides such as dichloro-diphenyl-trichloroethane (DDT) or dichloro-diphenyl-dichloroethylene (DDE), as a derivative of DDT, and the more toxic cyclodienes dieldrin and aldrin (Ratcliffe 1984; Wilson et al. 2018). The well-documented peregrine falcon population crash during the pesticide era showed a further reduction of the total UK breeding population by 50% within a decade (Ratcliffe 2003). By 1958, for example, there were no occupied territories and no young produced in Sussex, and peregrine falcons were absent from the region as a breeding species for over three decades (Franklin and Everitt 2009). The decline and subsequent crash of this southern England population of peregrines paralleled changes in the country as a whole (Ratcliffe 1984, 2003; Franklin and Everitt 2009). Therefore, during this time many peregrine breeding territories became vacant, acutely apparent across extensive areas of Southern England and Wales (Ratcliffe 1963; Banks et al. 2010). Only the peregrine populations of the Scottish Highlands, outer UK islands (Ratcliffe 1963) and Northern Ireland remained relatively unaffected (Wells and Ruddock 2009).

Since the ban on organochlorine pesticides in the UK in the 1980s and 1990s (Meijer et al. 2001), UK surveys of the peregrine breeding populations have recorded a steady increase (Ratcliffe 1984; Crick and Ratcliffe 1995; Horne and Fielding 2002; Wilson et al. 2018), and the recovery of the peregrine falcon is considered a flagship conservation success story (Crick and Ratcliffe 1995). In 2014, the breeding population estimates were 1769 pairs (Wilson et al. 2018), and by 2018, peregrine populations in England had increased by 250% of its former pre-war size (Wilson et al. 2018). National survey reports since 1972 document the progressive post-crash expansion of the peregrine populations notably in the north and west of the British Isles (Ratcliffe 1972, 1984; Crick and Ratcliffe 1995; Horne and Fielding 2002). However, the re-colonisation of Sussex by peregrine falcons lagged behind the rest of mainland UK, and it was not until 1990 when a single pair successfully bred (Franklin and Everitt 2009). In 2016, the Sussex peregrine population had substantially increased and was then composed of more than 40 breeding pairs, well in excess of the 8-12 pairs recorded between 1904 and 1954 (Franklin and Nicholls 2018). After an absence of more than 30 years, the remarkable and unaided recovery of the Sussex peregrine led conservation managers to question where the founders originated from, and if the current thriving population shares its ancestry with surviving native peregrines or with peregrines from further afield.

Such a newly formed population of peregrine falcons deriving from a few migrant founders could be experiencing detrimental impacts associated with inbreeding, genetic drift and low genetic variation (Maruyama and Fuerst 1984, 1985; Cornuet and Luikart 1996; Jacobsen et al. 2008). This could result in inbreeding depression (i.e. reduced biological fitness because of inbreeding) and deteriorated adaptive potential (Frankham 2015), including reduced egg clutch size and juvenile survival, often requiring intensive conservation intervention (Brown et al. 2007). As peregrine falcons have suffered unprecedented population bottlenecks and the occurrence of regional extirpations (Brown et al. 2007), it would be expected for the re-established Sussex peregrine population to be genetically impoverished, with limited potential for long-term survival. Conversely, levels of genetic diversity of a newly established population could be increased through mechanisms including hybridisation, mutation, and migration (gene flow; Hartl and Clark 2007).

As the origins of the Sussex peregrine population is unknown, it is therefore reasonable to hypothesise that the founder population could have been the result of immigration from these adjacent recovering populations, or even from peregrine populations in continental Europe to the east and Ireland from the west (Nicholls et al. 2017). There are, however, other plausible explanations to the rapid recovery of the Sussex peregrines. Fleming et al. (2011) speculated that losses of large numbers of domestic birds derived originally from captive-bred peregrines of several subspecies and even interspecific falcon hybrids could have augmented the recovering UK population. Extrapolation of Flemming et al. (2011) data and also information from a voluntary lost-found register for falconry birds (Barbara Royle pers. comm., The Independent Bird Register) indicate that on average around 30 peregrines and 30 peregrine hybrids (i.e. hybrids of peregrines with other *Falco* species) per year are reported as lost and unrecovered by mainland UK falconers since records began in 1980. This is likely an underestimate as there are no longer legal requirements to report losses or recoveries of falcons as there was previously. Furthermore, Flemming et al. (2011) speculated therefore that peregrines and peregrine hybrids have been inadvertently released in numbers which, based on planned re-introduction programmes, are known to be sufficient for establishing wild populations (Saar 1988; Holdroyd and Banash 1990; Tordoff and Redig 2001; Jacobsen et al. 2008). Moreover, a gyrfalcon/saker falcon x peregrine (of domestic origin) produced viable hybrid offspring with a wild peregrine within the Sussex population (Everitt and Franklin 2009). This key observation demonstrated that domestic hybrids between peregrine and other falcon species can interbreed and produce offspring with wild peregrines, and that domestic hybrids lost into the wild could potentially have a greater impact on the genetic diversity of the post-pesticide populations than the recruitment of mixed provenance domestic peregrines (*F. peregrinus ssp.*). Even a single pair which includes an escaped hybrid and a wild peregrine producing viable offspring in a strongly philopatric founder population could have a profound genetic effect on the allelic diversity of subsequent generations (Hartl and Clark 2007). The high phenotypic variation of Sussex peregrines noted by Nicholls et al. (2017) may provide further support for this hypothesis.

In this study we used a combination of mitochondrial (mtDNA) and microsatellite markers to examine the levels of genetic diversity and genetic structure for the population of peregrine falcons in the English south coast before their local extinction between 1955 and 1958 (pre-pesticide extinction) and since their repopulation of the area (post-pesticide recovery), to explain the origins of the recently recovered Sussex peregrines. For this, we analysed genetic data of: 1) a historic sample of peregrine falcons from museum specimens collected from the English south coast before the population crash of the mid 1950’s, 2) UK domestic-bred peregrines freely available for sale to falconers and therefore potentially vulnerable to loss into the wild population, 3) a German population which had an assisted recovery from local extinction by the release of very large numbers of captive-bred birds (Saar 1988; Wink 2019), 4) wild peregrines from the Republic of Ireland which had also experienced a population depletion in the 1960s and subsequent recovery (Ratcliffe 2003) but which has had no releases of captive-bred birds, and 5) a small sample of Mediterranean peregrines (*F.p. brookei*) which are a near geographical neighbour and thought to have contributed in some part to the recovery of the German peregrine population (Wink 2019).

Our aims were: 1) to assess the levels of genetic diversity in the re-established Sussex peregrine population compared with a historical population of peregrine falcons before the population decline; 2) to test for signatures of genetic bottlenecks in the contemporary population; 3) to estimate the genetic structure among British, Irish, continental European and domestic-bred peregrine falcons, as well as between contemporary and historical populations; and 4) to evaluate the potential phylogeographic origin of the Sussex peregrines from wild peregrines or from augmentation by domestic peregrines of mixed origin.

## Methods

### Sample collection and groupings

A total of 164 feather or toe-pad samples were obtained from various regions (Online Resource 1 and Table S1). Contemporary feather samples of peregrine falcons (*F. p. peregrinus*) from the south of England were collected from wild birds by the Sussex Peregrine Study over the period of 2000 to 2015 from known peregrine territories and monitored nesting sites within the County of Sussex, and from London, Hampshire, Kent and Surrey (N = 37). Feather samples were obtained from Ireland (wild *F. p. peregrinus* or first generation bred from wild birds obtained in the Republic of Ireland, N = 30) where there was a population decline of peregrines in the 1960’s followed by a rapid, and probably unassisted, re-establishment (Ratcliffe 1984; Norriss 1995; Madden et al. 2009) with a recent census of 425 pairs in 2017 (David Norriss pers. comm.). Feather samples were obtained from North-Rhine Westphalia in Germany (wild free-living peregrine falcons, N = 44). In Germany, peregrines were extirpated during the 1950’s and 60’s with only around 40 pairs remaining, these in the south of the country (Saar 1988) where there was an assisted reintroduction of just over 1000 captive bred birds comprising some *F. p. brookei* but predominantly *F. p. peregrinus* (Saar 1988; Wink 2019). Feathers samples were taken from randomly selected domestic birds from different breeding lines from various breeders (‘peregrines’ bred from a mix of races and subspecies of *F. peregrinus* and sibling species, including *F. p. pelegrionides* and *F. p. babylonicus*, of largely unknown provenance; N = 30). The founder breeding population for domestic birds was around 200-250 breeding birds and of these approximately 2/3 were native *F. p. peregrinus* and 1/3 imported birds of other subspecies (Online Resource 1). Other feather samples were obtained from Mediterranean peregrines (*F. p. brookei*) from Spain, Gibraltar and Italy (N = 5). The relative proximity of Mediterranean peregrines to *F. p. peregrinus* populations in UK and mainland Europe makes it a potential candidate as a source of immigrants into depleted post-pesticide *F. p. peregrinus* populations. Toe-pad samples of peregrine falcons from British museums (N = 18) were obtained from several counties in the south of England and the Channel Islands (Table S1). For this study, museum samples dated pre-1955 represented the pre-pesticide population crash, while toe-pad and feather samples with a collection date post-1955 represented the post-pesticide recovery. Samples were stored at 4 °C or at room temperature until analysis.

The total sample was then divided into geographical and temporal groups for further analysis with microsatellite and mtDNA data, namely: ‘UK-post’ (UK post-1955 ban on organochlorine pesticides, including museum samples dated post-1955, feathers from wild free living animals, and wild-injured animals in captivity), ‘UK-pre’ (UK pre-1955 ban on organochlorine pesticides, including museum samples pre-1955), ‘Domestic’ (all post-1955 feathers from animals bred in captivity in the UK, as explained above), ‘German’ (all post-1955 feathers from wild free-living animals), ‘Irish’ (all post-1955 feathers from wild, wild-injured birds in captivity and F1 from wild-take) and *‘F. p. brookei*’ (including feathers from wild-injured animals held in captivity for rehabilitation and captive bred). To explore the genetic polymorphism at the UK and at the species levels, the groups ‘UK-peregrines’ (including all pre- and post-1955 samples) and ‘All-peregrines’ (all *F. peregrinus* and subspecies from all sources) were also created.

### DNA extraction

DNA from contemporary feathers was extracted using the GeneJET Genomic DNA Purification Kit (ThermoFisher Scientific) or the using Bioline Isolate II Genomic DNA extraction kit (Bioline) following the manufacturer’s protocol and suspended in 50 μL elution buffer. DNA from historical museum toepad samples was extracted using the Bioline Isolate II Genomic DNA extraction kit (Bioline), but suspending tissues in 400 μL lysis buffer with 25 μL proteinase k and incubated overnight at 55 °C. DNA was then washed through a spin column and eluted into 50 μL elution buffer. DNA extractions from historical museum toepad samples were carried out in a separate laboratory dedicated to ancient DNA work and in a UV-irradiated fume hood to ensure no contamination. Negative controls were included to ensure no contamination.

### Microsatellite genotyping

We used a suite of ten polymorphic microsatellite markers developed for peregrine falcons (NVHfp5, NVHfp13, NVHfp31, NVHfp46-1, NVHfp54, NVHfp79-4, NVHfp82-2, NVHfp86-2, NVHfp89, NVHfp92-1; Nesje et al. 2000). The PCR procedure and amplification volumes followed Raisin et al. (2009) for all samples (feather and toe-pad), set at a total volume of 2 μL. Each PCR contained 2 μL of DNA that was air-dried, 1 μL of primer mix (with a fluorescently labelled forward primer) at 0.2 μM and 1 μL QIAGEN Multiplex PCR Master Mix (QIAGEN). PCR cycling conditions were 95 °C for 10 min followed by 35 cycles of 95 °C for 30 s, 55 °C for 90 s and 72 °C for 90 s, with a final incubation at 72 °C for 10 min. PCR products were separated using an Applied Biosystems 3730 DNA analyser (Applied Biosystems), using ROX 500 as a size standard. Alleles were scored on GeneMapper version 3.7 (Applied Biosystems).

Approximately 25% of contemporary samples were amplified twice at all 10 loci to determine genotyping error (Hoffman and Amos 2005; Pompanon et al. 2005), and 37.5% of museum samples were genotyped twice to confirm accurate scoring of alleles and to allow identification of potential errors due to allelic dropout of larger alleles. Owing to degradation and yield associated with historical DNA (Wandeler 2007), two microsatellite markers, NVHfp5 and NHVfp92-1, failed to amplify across all museum samples. Genotype error (mismatch alleles) was found for microsatellite marker NHVfp82-2 across all DNA samples and was subsequently removed from analysis. The resulting dataset comprised seven microsatellite markers and 149 individuals (available on Dryad https://doi.org/10.5061/dryad.mgqnk98wj).

### Microsatellite analysis

Any potential genotyping errors due to microsatellite stuttering, allelic dropout and null alleles in the total sample were examined using Micro-Checker version 2.2.3 (Van Oosterhout et al. 2004). Hardy-Weinberg equilibrium (HWE) and linkage disequilibrium (LD) between loci were calculated using Genepop version 4.6 (Raymond and Rousset 1995; Rousset 2008).

To assess the levels of genetic diversity of peregrine falcons, the allelic richness (i.e. number of alleles, N_a_), effective number of alleles (N_e_), observed heterozygosity (H_O_), and measured heterozygosity (H_e_; Nei’s gene diversity), inbreeding coefficient (F_IS_), and private alleles (N_p_) per population were estimated in GenAlEx version 6.5 (Peakall and Smouse 2012). A Multivariate Analysis of Variance (MANOVA) on the per-locus genetic diversity values (N_a_, N_e_, H_O_ and H_e_) was done in SPSS version 24 (SPSS Statistics) to determine whether there were temporal differences between UK-post and UK-pre populations. The effective number of breeders (N_eb_), or inbreeding effective number, and the jackknife confidence intervals were estimated for each population using the co-ancestry method of Nomura (2008), as implemented in NeEstimator version 2.01 (Do et al. 2014). The presence of genetic bottlenecks was tested by estimating heterozygosity excess based on the two-phase model (TPM; 70% proportion of stepwise mutation model and variance = 30) and the infinite-alleles model (IAM) as implemented in Bottleneck version 1.2 (Piry et al. 1999). In populations undergoing bottleneck, the allelic richness is reduced faster than heterozygosity (Nei’s gene diversity) thus becoming larger than the expected heterozygosity at mutation-drift equilibrium (Cornuet and Luikart 1996), and the difference was tested with a Wilcoxon’s test assuming that all loci fit the TPM or IAM under mutation-drift equilibrium.

Nei’s unbiased pairwise genetic distances (Nei 1978) were estimated among populations in GenAlEx which for neutral markers under an infinite-allele-model it is predicted to increase linearly with time. The extent of population subdivision was assessed by calculating Wright’s pairwise differentiation values among populations (F_ST_), and by a hierarchical analysis of molecular variance (AMOVA) at three hierarchical levels (among populations, among individuals within populations and within individuals) using GenAlEx. Significance levels were obtained using 1000 permutations.

Genetic distances among individuals were estimated (Smouse and Peakall 1999), and the distance matrix was converted to a covariance matrix to perform a Principal Coordinate Analysis (PCoA) in GenAlEx. The detection of first-generation migrants was done with GeneClass2 (Piry et al. 2004) selecting likelihood ratio L_home_/L_max_, the Bayesian method by Rannala and Mountain (1997), 1000 Markov Chain Monte Carlo simulations (Paetkau et al. 2004), and a probability threshold alpha = 0.05. The assignment of individuals to populations (using the leave-one-out procedure) based on their relative likelihood scores and a quality index (computed as the mean value of the scores of each individual in the population it belongs to) were also estimated with GeneClass2. A Bayesian approach in Structure version 2.3.4 (Pritchard et al. 2000) was used to detect the most likely number of genetic clusters among peregrine falcon populations doing 10 replications with number of clusters K = 1–10 with 100,000 burn-in, 1,000,000 MCMC iterations after burn-in, and admixture model (using sampling locations as prior information) with correlated allele frequencies (Falush et al. 2003). The most likely number of K clusters was examined in StructureSelector (Li and Liu 2018) using log likelihood scores [mean LnP(K)] and ΔK values (Evanno et al. 2005). As these methods often underestimate clusters due to uneven sample sizes (Janes et al. 2017), we obtained estimates of K based on Puechmaille’s method by subsampling the independent clusters previously identified, a technique which accounts for uneven sample size across groups (Puechmaille 2016). Likelihood scores and clusters were obtained using the CLUMPAK (Kopelman et al. 2015) function in StructureSelector and individual probability plots were generated using Structure Plot (Ramasamy et al. 2014).

### Mitochondrial DNA sequencing

PCR was performed to amplify approximately 650 bp of the domain I of the mtDNA control region of contemporary peregrine falcon feathers using primers L15206 and H15856 (Talbot et al. 2011). PCR amplifications were carried out in 50 μL final volume, consisting of 5 μL 10X Maxima Hot Start Taq buffer, 5 μL dNTP Mix (2 mM each), 1 μM 10μM forward and reverse primers, 1.2 μL 25 mM MgCl_2_, 0.3 μL 5U/μL Maxima Hot Start Taq DNA Polymerase (ThermoFisher Scientific), 1 μL DNA sample and 27.5 μL molecular grade water. PCR conditions were 95° C for 5 min followed by 35 cycles of 95° C for 60 s, 47° C for 60 s and 72° C for 90 s, and a final elongation step at 72° C for 10 min. PCR products were visualised under UV light after electrophoresis in 1% Agarose gels containing SYBR Safe DNA gel stain (Invitrogen). Successful amplifications were purified using the GeneJET PCR Purification Kit (ThermoFisher Scientific) and send for Sanger sequencing to DBS Genomics (Durham University).

PCR amplification for museum toepad samples was carried out in a separate irradiation UV isolation hood dedicated to ancient DNA work to prevent contamination. Due to the fragmented nature of museum DNA, internal primers were specifically designed to amplify short overlapping fragments (175-198 bp) of the mtDNA control region of peregrine falcons (Table S2). PCR amplifications were carried out in 10 μL final volume, consisting of 5 μL MyTaq HS DNA Polymerase (Bioline), 0.2 μL of each forward and reverse primer, 1 μL DNA and 3.6 μL dH_2_O. Negative controls were included in each PCR replacing DNA with ultrapure ddH_2_O. PCR cycling parameters were an initial hot start of 95 °C for 1 min, followed by 35 cycles of 95 °C for 15 s, 52 °C for 15 s, 72 °C for 10 s, followed by a final incubation period of 72 °C for 10 min. PCR products and negative controls were examined by agarose gel electrophoresis to ascertain amplification and to confirm no traces of contamination. PCR product was purified and sequenced using a 3730xl analyser (Macrogen).

From the feather and toepad samples, 102 DNA sequences were newly obtained here, from which 100 were *F. peregrinus* and two were *F. rusticolus*, and all new sequences were deposited in GenBank (accession numbers: MT247108-MT247209). In addition, 83 DNA sequences of peregrine falcons (including sequences identified to the species level as *F. peregrinus*, and to the subspecies level *F. p. peregrinus* or other subspecies of peregrines) were retrieved from GenBank (accession numbers: AF090338, DQ165502-DQ165549, JN400833-JN400846, JQ282801, JX029991, JX878245-JX878269, and NC000878), plus eight gyrfalcons *F. rusticolus* (accession numbers: EF517337-EF517344) used as an outgroup.

### Mitochondrial DNA analysis

All control region sequences were aligned using ClustalW multiple alignment in BioEdit version 7.2.6 (Hall 1999). Considering alignment gaps, 628 bp were analysed in a total of 193 individuals. DNA sequences were imported into DnaSP version 5.10.1 (Librado and Rozas 2009) and were separated into the main groups indicated above (i.e. UK-post, UK-pre, German, Domestic, Irish and *F. p. brookei*), and into ‘GenBank-peregrines’ (identified from online source as the subspecies *F. p. peregrinus*), and ‘GenBank-others’ (identified from online source as the species *F. peregrinus* and/or several other subspecies of peregrines but not *F. p. peregrinus* or *F. p. brookei*), and ‘*F. rusticolus*’ (i.e. gyrfalcons, the outgroup to peregrine falcons; identified from online source, and feathers from captive bred animals).

DNA polymorphism indices were obtained in DnaSP to allow comparisons among the groups and published data, including total number of segregating sites (S), number of polymorphic sites (Pol), number of haplotypes (H), haplotype diversity (Hd), nucleotide diversity (π), and average number of nucleotide differences (k). To explore the historical demography of the falcons, pairwise sequence polymorphisms were estimated to generate mismatch distributions, and the population size changes per group were evaluated using Ramos-Onsins and Rozas’ R_2_ (Ramos-Onsins and Rozas 2002) and Fu’s Fs (Fu 1997) values calculated with DnaSP, with significance tested with the coalescent process for a neutral infinite-sites model and assuming a large constant population size (1000 replications).

Pairwise genetic differentiation (F_ST_) among groups were obtained using DnaSP to analyse the genetic structure of the populations. An AMOVA (among and within populations) was performed using Arlequin version 3.5 (Excoffier and Lischer 2010), with significance levels were obtained using 1000 permutations. The phylogenetic relationships among the samples was assessed with a phylogenetic network using all the haplotypes, including those from museum samples, generated with the software Network version 5.0.0.3 (Fluxus Technology). The Median Joining algorithm with the Greedy FHP criterion was selected.

## Results

### Microsatellite data analysis

There was no evidence of large allele dropout. Only one locus showed potential scoring errors due to stuttering but revising the genotype data showed no allele scoring errors. Four out of seven loci could have null alleles, but this was probably due to homozygosity excess for most allele size classes. Overall, estimated frequencies of null alleles for the full dataset ranged between 0.010%–0.15%, which was considered acceptable due to the origin of the feather and tissue samples. There were no deviations from HWE in the populations due to heterozygote excess or in a global test across all loci across all populations; however, deviations from HWE due to heterozygote deficiency were detected in four populations (UK-post, Domestic, German and Irish, P < 0.05). There was no evidence of LD for each locus pair across all populations.

Overall, genetic diversity values were lower in the UK-post and UK-pre groups compared with other peregrine falcon groups, and the UK-post group had the lowest heterozygosity value and number of effective alleles (N_e_) among all populations (Table 1). Comparing the two temporal groups, UK-post had higher mean number of alleles (N_a_) and heterozygosity (H_e_) than UK-pre, but lower observed heterozygosity (H_O_) and number of private alleles (N_p_), the Domestic group had the highest N_p_ followed by the Irish group, and the *F. p. brookei* group had the highest level of heterozygosity followed by the Domestic and Irish groups (Table 1; Online Resource 1 and Fig S1). However, the differences in genetic diversity values were not statistically significant (MANOVA: Wilk’s λ F = 0.933, P = 0.487; Box’s test of equality of covariances: F = 1.834, P = 0.052; Levene’s tests of equality of variances: P > 0.05 for all variables). The Domestic group had the highest value of inbreeding (F_IS_) while the *F. p. brookei* and UK-pre groups had the lowest levels of inbreeding (Table 1). The UK-post and German groups had the lowest effective number of breeders (N_eb_), while the Domestic group had the highest N_eb_ (Table 1). There was no evidence of genetic bottleneck under the TPM with all P-values > 0.05; however, the UK-post (P = 0.039), UK-pre (P = 0.016) and German (P = 0.039) groups showed signatures of genetic bottlenecks under the IAM. There were significant results for the *F. p. brookei* group under the TPM and IAM, but caution should be taken because of low sample size.

**Table 1.** Genetic diversity values in peregrine falcon (*Falco peregrinus*) populations based on microsatellite data

|  |  |  |  |  |  |  |  |  |  |  |  |  |
| --- | --- | --- | --- | --- | --- | --- | --- | --- | --- | --- | --- | --- |
| Population | N | N<sub>miss</sub> | N<sub>a</sub> | N<sub>e</sub> | H<sub>o</sub> | H<sub>e</sub> | F<sub>IS</sub> | N<sub>P</sub> | N<sub>P</sub><br>(prop.) | N<sub>eb</sub> (CI) | N<sub>assign</sub> | %<br>Assign |
| UK-post | 39 | 35.143 | 4.143 | 2.294 | 0.557 | 0.548 | -0.005 | 2 | 0.286 | 4.5 (1.4-9.6) | 30 | 76.923 |
| UK-pre | 14 | 8.571 | 3.286 | 2.297 | 0.587 | 0.545 | -0.097 | 3 | 0.429 | Infinite | 8 | 57.143 |
| German | 30 | 22.143 | 4.429 | 2.649 | 0.581 | 0.590 | 0.008 | 1 | 0.143 | 6.5 (2.9-11.6) | 24 | 88.889 |
| Irish | 30 | 23.143 | 6.000 | 3.481 | 0.634 | 0.669 | 0.029 | 3 | 0.429 | 19.3 (0.5-71.3) | 21 | 72.414 |
| Domestic | 30 | 19.857 | 7.000 | 4.122 | 0.624 | 0.706 | 0.114 | 10 | 1.429 | 81.7 (0.1-410) | 19 | 65.517 |
| *F. p. brookei* | 6 | 4.714 | 3.714 | 2.885 | 0.771 | 0.646 | -0.194 | 0 | 0.000 | 30.9 (0-155.4) | 4 | 66.667 |
| All UK-peregrines | 53 | 43.714 | 4.714 | 2.302 | 0.558 | 0.558 | 0.003 | 6 | 0.857 | 22.3 (0-111.7) | 52 | 98.113 |
| All-peregrines | 149 | 113.571 | 9.429 | 3.818 | 0.599 | 0.725 | 0.170 | - | - | 10.3 (5-17.6) | - | - |
| N = sample size, N<sub>miss</sub> = sample size considering missing data across loci, N<sub>a</sub> = number of different alleles, N<sub>e</sub> = number of effective alleles, H<sub>o</sub> = observed heterozygosity, H<sub>e</sub> = expected heterozygosity, F<sub>IS</sub> = fixation index, N<sub>P</sub> = number of private alleles, N<sub>P</sub> (prop.) = proportion of private alleles in the population, N<sub>eb</sub> = Effective number of breeders (CI = jackknife confidence interval), N<sub>assign</sub> = Number of individuals used in assignment test, % Assign = percentage of individuals correctly assigned to the population of origin |  |  |  |  |  |  |  |  |  |  |  |  |

There were high pairwise genetic distances between the UK-post and UK-pre groups with any of the other groups, while the lowest genetic distance among all pairwise comparisons was between the German and *F. p. brookei* groups, and between the UK-post and UK-pre groups (Table S3). All pairwise comparisons (except between the German and *F. p. brookei* groups) showed significant and moderate levels of genetic differentiation (F_ST_) (Table S3); the highest F_ST_ values were observed in pairwise comparisons between the UK-post and UK-pre groups with any other groups (Table S3). Based on the AMOVA, 13% of the total variation was found among groups, while 30 and 57% was found among and within individuals, respectively, and significant global F-statistics (F_ST_ = 0.130, F_IS_ = 0.348 and F_IT_ = 0.433, all P < 0.05).

A total of 37.9% accounted for the first three dimensions of the PCoA (PCo1 = 19.28%, PCo2 = 10.02% and PCo3 = 8.6%). The analysis indicated two main clusters, one including the UK-post and UK-pre samples, and another one mostly including all other samples (Fig. 1). Eight individuals were detected as first-generation migrants with likelihood probabilities below 0.05, including four from UK-post assigned to UK-pre, one from UK-post assigned to the German group, one from the Domestic group and from *F. p. brookei* assigned to the Irish group, and one from *F. p. brookei* assigned to the German group. Based on the genotypes, 73.6% of the individuals (106 out of 144 individuals with sufficient genotype information) were correctly assigned to the group of origin (quality index = 68.65%); the German group had the highest percentage of correct assignment of individuals (88.9%), followed by UK-post (76.9%), Irish (72.4%), *F. p. brookei* (66.7%), Domestic (65.5%) and UK-pre (57.1%). Structure analysis using sub-sampling using Puechmaille’s method resulted in the most likely number of clusters at K = 4 (Fig. 2). Assuming K = 4, the UK-pre and UK-post individuals assigned predominantly to cluster 1, whilst Domestic individuals assigned to cluster 2, German and *F. p. brookei* individuals predominantly assigned to cluster 3, and Irish individuals assigned to cluster 4 (Fig. 2). However, the log likelihood simulations indicated K = 2 clusters under Evanno’s method, while mean LnP(K) indicated K = 3 clusters. Assuming K = 2 or K = 3, the UK-pre and UK-post peregrines also formed one cluster, demonstrating the genotypic similarities of the two temporal groups (Fig. 2).

**Fig. 1.**
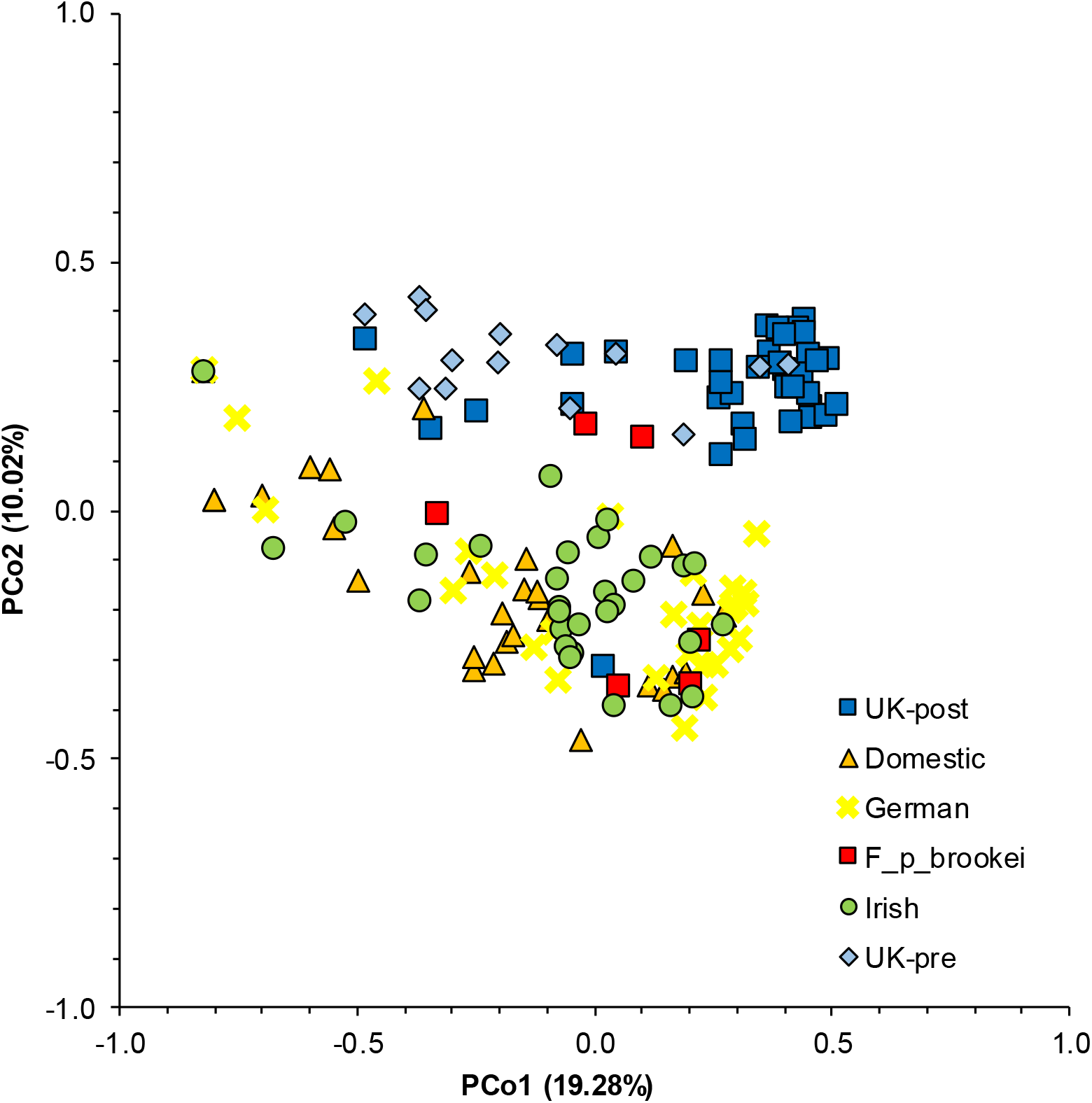
Principal coordinates analysis (PCoA) based on microsatellite genotypes of peregrine falcons (*Falco peregrinus*)

**Fig. 2.**
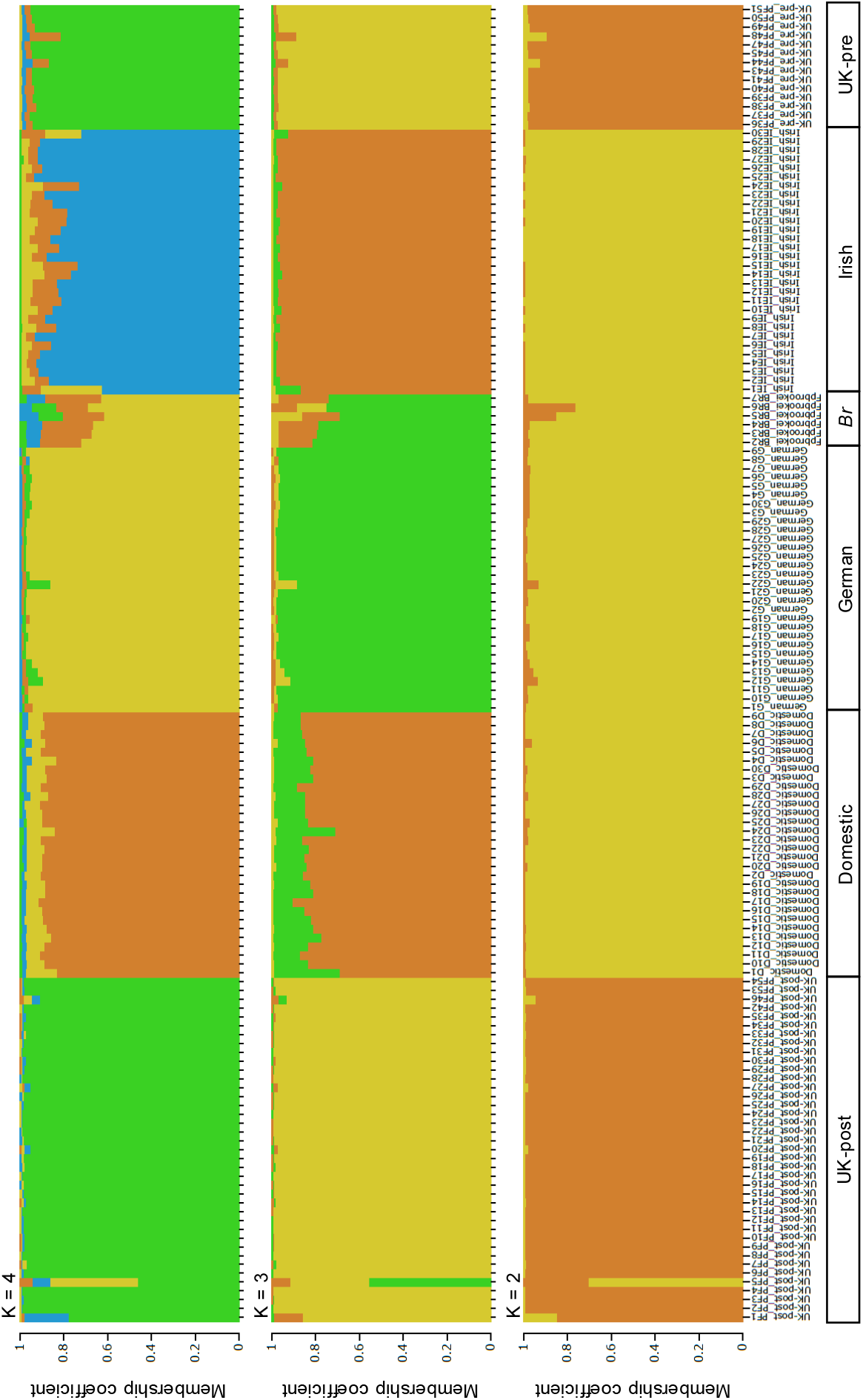
Individual probability plots showing the most likely number of genetic clusters of peregrine falcons (*Falco peregrinus*) for K number of clusters (K = 4, 3 and 2) based on a Bayesian analysis of genetic structure of microsatellite data

### Mitochondrial DNA data analysis

A 628 bp fragment of the control region was obtained for all feather samples from contemporary birds. However, probably because of DNA degradation and amplification failure of 5’ and/or 3’ regions, it was not possible to obtain the full mtDNA control region fragment from some museum (toepad) samples (pre- and post-1955). This restricted the amount of nucleotide sites available for pairwise sequence comparisons when using museum samples in UK-pre and UK-post groups.

A total of 183 mtDNA sequences of peregrine falcons were obtained for analysis from all sources (including GenBank, museum and feather samples), but only 10 haplotypes were found (Table 2). Haplotype diversity varied considerably among peregrine groups, with the lowest values found in the UK-pre and UK-post groups, and the highest values found in the Irish group, followed by the Domestic, GenBank-peregrines and German groups (Table 2). Nucleotide polymorphism was also generally low across all peregrine groups, with the lowest values observed in German and UK-post groups (Table 2).

**Table 2.** DNA polymorphism in peregrine falcon (*Falco peregrinus*) populations based on mtDNA control region data

|  |  |  |  |  |  |  |  |  |  |  |
| --- | --- | --- | --- | --- | --- | --- | --- | --- | --- | --- |
| Population | N | S | Pol | H | Hd | $\pi$ | k | $R_2$ | $F_s$ | $\tau$ |
| UK-post | 24 | 363 | 6 | 5 | 0.529 | 0.002 | 0.837 | 0.101 | -1.349 | 0.356 |
| UK-pre | 14 | 295 | 9 | 5 | 0.505 | 0.005 | 1.407 | 0.132 | -0.786 | 0.153 |
| German | 21 | 627 | 2 | 4 | 0.710 | 0.001 | 0.924 | 0.231 | -0.140 | 0.924 |
| Irish | 22 | 627 | 5 | 8 | 0.892 | 0.003 | 1.792 | 0.179 | -2.349 | 1.792 |
| Domestic | 14 | 627 | 4 | 5 | 0.824 | 0.002 | 1.209 | 0.148 | -1.159 | 1.209 |
| GenBank-peregrines | 55 | 376 | 6 | 7 | 0.756 | 0.003 | 1.182 | 0.098 | -1.289 | 1.182 |
| GenBank-others | 28 | 556 | 26 | 19 | 0.971 | 0.006 | 3.198 | 0.087 | **-13.598** | 0.828 |
| *F. p. brookei* | 5 | 627 | 2 | 3 | 0.800 | 0.002 | 1.000 | 0.250 | -0.475 | 1.000 |
| UK-peregrines | 38 | 290 | 11 | 8 | 0.486 | 0.003 | 0.908 | 0.070 | **-3.908** | 0.095 |
| All-peregrines | 183 | 238 | 11 | 10 | 0.320 | 0.002 | 0.365 | 0.026 | **-9.361** | 0.365 |
| *F. rusticolus* | 10 | 627 | 8 | 7 | 0.911 | 0.003 | 1.867 | **0.116** | **-3.502** | 1.867 |
| N = number of sequences used, S = total number of sites (excluding sites with gaps/missing data, or excluding them in pairwise comparisons), Pol = polymorphic sites, H = number of haplotypes, Hd = haplotype diversity, $\pi$ = nucleotide diversity, k = average number of nucleotide differences, $F_s$ = $F_u$ (1997), $R_2$ = Ramos-Onsins and Rozas (2002), $\tau$ = 2ut. Values in bold ($P < 0.05$), simulations based on the coalescent process for neutral infinite-sites model assuming a large constant population size | | | | | | | | | | |

No signatures of a recent population expansion were detected in any peregrine falcon group with R_2_ or Fs (Table 2); however, *F. rusticolus* showed significant departures from neutrality using R_2_. Based on Fs values, the groups GenBank-others, UK-peregrines, All-peregrines and *F. rusticolus* showed significant departures from neutrality. Overall, there were very few pairwise differences among mtDNA sequences of peregrine falcons and all mismatch distributions were right skewed (Fig. 3); UK-pre and UK-post groups showed high frequencies of 0 pairwise differences, and other peregrine falcon groups showed high frequencies of only 1-2 pairwise differences.

**Fig. 3.**
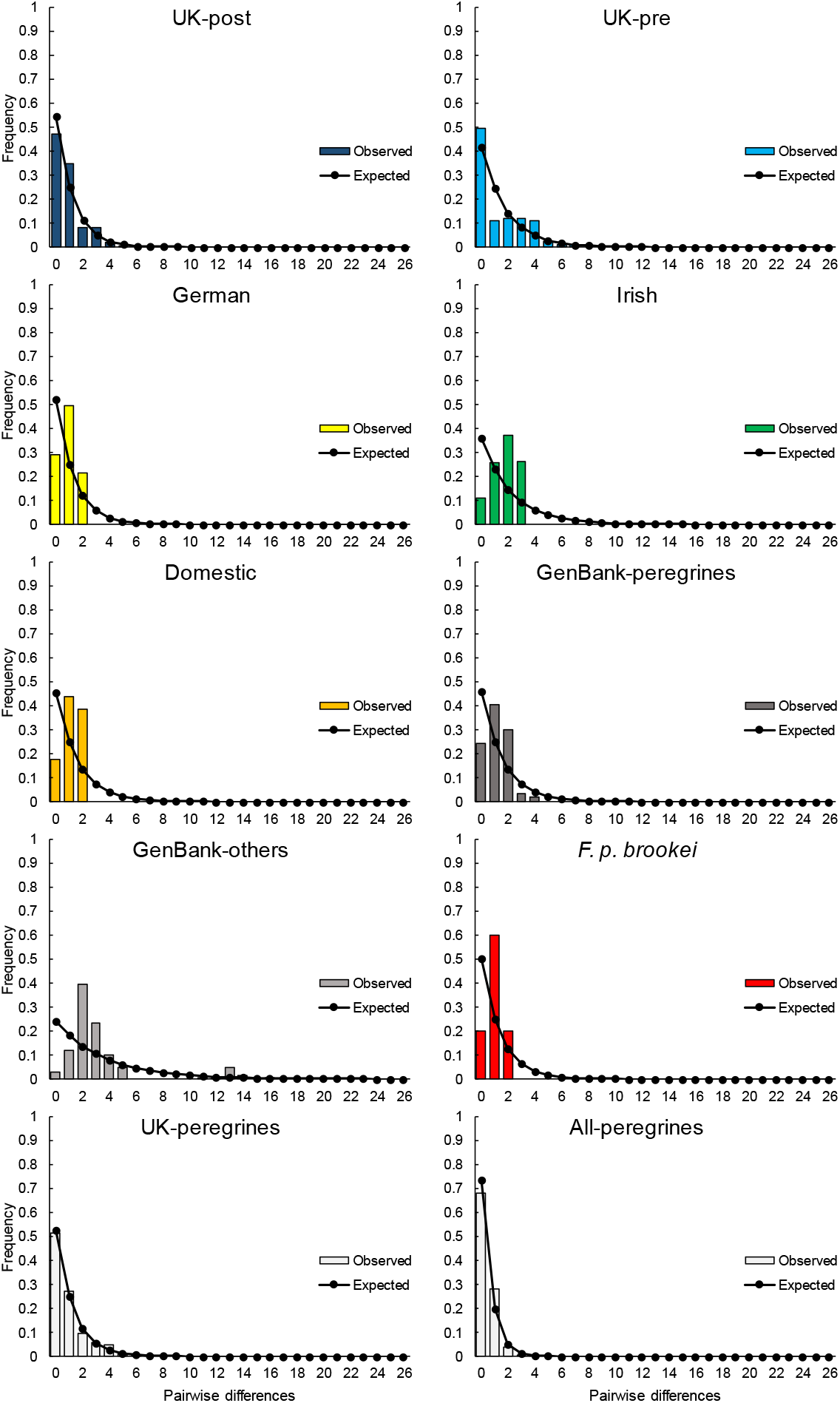
Mismatch distributions of peregrine falcon (*Falco peregrinus*) populations showing the observed pairwise differences in mtDNA control region and expected values under neutrality

Pairwise genetic differentiation (Table S4) among all peregrine falcon groups (i.e. excluding *F. rusticolus*) showed a low average F_ST_ = 0.064 (or 6.4% differentiation in the mtDNA control region), with values ranging from 0 to 20.4%, indicating low genetic differentiation. There was no differentiation (0%) between UK-post vs. Domestic, Irish vs. GenBank-others, and German vs. *F. p. brookei* groups, very low differentiation between UK-post vs. Irish groups (0.8%), and low differentiation between UK-post vs. UK-pre groups (5%). Higher differentiation values were obtained in pairwise comparisons with GenBank, GenBank-others, *F. p. brookei* and *F. rusticolus* groups, as expected from their taxonomic diversity or evolutionary divergence. Based on the AMOVA, 43.86% and 56.14% of the total variation was found among and within groups, respectively, and there was a significant global FST = 0.439 (P < 0.05).

The phylogenetic haplotype network (Fig. 4) reflected the genetic diversity and differentiation results shown above. Including museum samples increased the number of sites with gaps/missing data which cannot be utilised in phylogenetic haplotype network reconstructions; therefore, the network analysis was based on 238 nucleotide sites. There were 13 haplotypes in the total sample, all peregrine samples belonged to 10 haplotypes, while all *F. rusticolus* samples (the outgroup) belong to three other haplotypes. Most peregrine falcon samples belonged to haplotype 1 (150 samples), which had a central position relative to all other peregrine haplotypes, and it was formed by a mix of samples from various peregrine groups, including 10 UK-pre, 17 UK-post, 21 German, 20 Irish, 10 Domestic, 45 GenBank-peregrines, 22 GenBank-others and five *F. p. brookei*. Other peregrine samples (15) belonged to haplotype 2, which included samples from all peregrine falcon groups except UK-pre, German and *F. p. brookei*. Overall, the group GenBank-others showed the highest number of haplotypes, including three haplotypes not found elsewhere, reflecting world-wide origin of the GenBank data set. The German group was the least diverse with only one haplotype, while the Irish and Domestic groups only had two haplotypes. UK-post and UK-pre had four haplotypes both; notably, UK-pre was represented in four haplotypes, two shared with UK-post and two unique haplotypes (haplotypes 5 and 6), each separated from the central haplotype 1 by two mutational steps; UK-post had only one unique haplotype (haplotype 7) not shared with any other group.

**Fig. 4.**
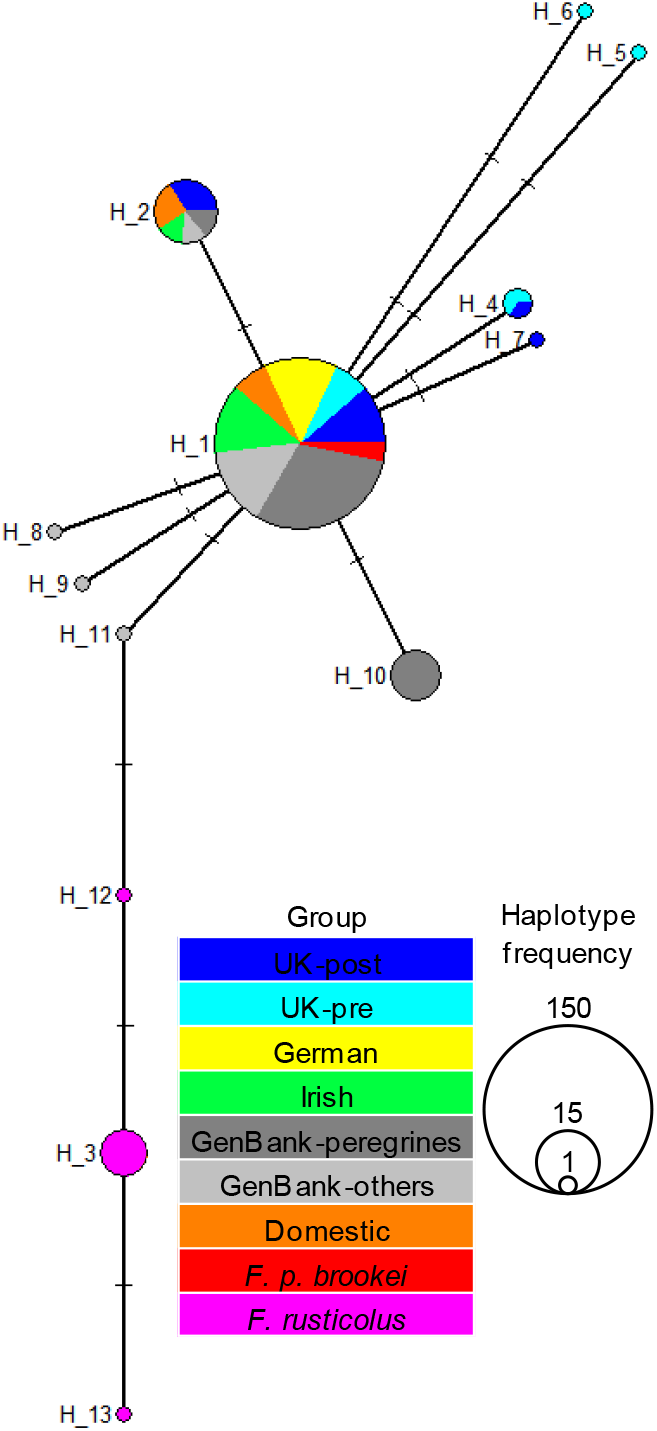
Phylogenetic haplotype network showing the evolutionary relationships among peregrine falcon (*Falco peregrinus*) mtDNA control region haplotypes arranged into groups

## Discussion

Using microsatellite and mtDNA markers, we examined the levels of genetic diversity and genetic structure of peregrine falcon populations before and after the documented post-pesticide extinction in the English south coast, compared the genetic affinities of UK peregrines with other European populations, domestic-bred birds of unknown provenance and with published falcon mtDNA data, and explored the phylogeographic origins of the recently recovered Sussex peregrines. The genetic data was obtained from museum specimens collected from the English south coast before and after the population crash of the mid 1950’s, from contemporary UK wild and captive specimens, and from contemporary Irish and continental European wild and captive specimens, and UK domestic-bred peregrines of mixed ancestry. Here, we report the unassisted recovery of a peregrine falcon population in Southern England with no detected signature of a genetic bottleneck and little change in the level of genetic diversity through time. This adds to the growing literature showing unassisted recoveries of peregrines in several countries (Sielicki and Mizera 2009) and, to our knowledge, this is the first study to explore this in the UK using molecular tools, with implications toward their conservation and protection.

### Spatial and temporal genetic diversity and structure

Successful breeding, good dispersal capabilities and inter-population connectivity are fundamental for preventing inbreeding depression and subsequent loss of genetic diversity (Frankham 2015; Ponnikas et al. 2017). As evidenced by our population genetics and phylogeographic study, results broadly indicated that there was no overall loss of genetic diversity or signature of a population bottleneck, although some genetic diversity indices showed lower genetic variation in the post-pesticide population (UK-post) than in the pre-pesticide population (UK-pre). Based on microsatellite data, there was, however, a difference in the number and proportion of private alleles, with UK-post showing lower values than UK-pre, and the UK-pre and UK-post were less differentiated genetically with each other than with any other peregrine group. Based on mtDNA, the UK post-pesticide population also showed lower number of polymorphic sites and nucleotide diversity values than the pre-pesticide population. UK-pre had two haplotypes not found elsewhere in our samples, which could represent a significant loss of haplotype diversity; however, there was very low genetic differentiation in mtDNA control region overall, as previously described in cytochrome b in peregrine falcons (Wink 2019).

It was expected, based on population genetics theory, that the re-established Sussex peregrine population would be genetically impoverished due to being founded by a small number of individuals surviving the pesticide crash. It was therefore somewhat surprising the level of genetic diversity retained by the newly established population, comparable to ancestral levels in the UK, and also remarkable considering that no specific conservation efforts (i.e. no formal reintroductions) have been directed to peregrines in the UK, a conservation strategy that other countries have employed (Brown et al. 2007; Jacobsen et al. 2008; Johnson et al. 2010). We have observed no genetic evidence of informal reintroductions via falconry losses of domestic birds.

The results found here are consistent with other spatial and temporal studies on falcons using microsatellites and/or mtDNA control region. In general, it would appear that the heterozygosity levels have remained comparable among peregrine falcon populations from the UK and other regions of the world in spite of the presumed relatively recent population-bottleneck, and there also appears to be differences in allelic diversity, with the UK (pre- and post-pesticide), Irish and German populations having fewer alleles than other peregrine falcon populations, but the Domestic group showing higher levels.

Compared with this study, for example, using a panel of 11 polymorphic microsatellite loci Brown et al. (2007) showed similar levels of heterozygosity (but slightly higher allelic richness) in historical and contemporary Canadian peregrine falcon (*F. p. tundrius*) populations, no signatures of genetic bottlenecks, and weak but significant population genetic structure; however, effective number of breeders in the Canadian falcon populations were much higher than those reported here. Based on mtDNA, Brown et al. (2007) found low nucleotide and haplotype diversities in historical and contemporary populations, much lower than in the UK-post and UK-pre populations reported here. Overall, Brown et al. (2007) concluded that contemporary and historical samples had similar genetic diversity levels, and that the organochlorine-induced bottleneck did not likely had an impact on the evolutionary potential of peregrines in Canada.

While studying captive and wild populations of peregrines in Scandinavia and potential gene introgression from the captive breeding stock during a reintroduction programme using 11 microsatellite loci, Jacobsen et al. (2008) showed similar heterozygosity values among contemporary, captive and historical populations, which were comparable with the heterozygosity values reported here; however, they found higher allelic richness than in the wild populations of peregrines studied here. Jacobsen et al. (2008) therefore concluded that the levels of genetic diversity in the contemporary wild population of peregrines in Scandinavia remained higher than expected from the population bottlenecks, and that this could be due to population admixture, considerable gene introgression and the release of a large number of young from the captive breeding population, and increased survivorship of chicks and adults in the wild, which could have altered the genetic composition of the contemporary wild population.

Ponnikas et al. (2017), using 10 microsatellite loci, also found similar heterozygosity values and no genetic bottlenecks, but they found higher allelic richness in peregrine falcons from Finland than in the UK populations reported here, and no genetic structure. They concluded that the genetic diversity could have been maintained through high dispersal rates across Finland without reintroductions, thus preventing genetic structuring of populations and the negative effects of the population decline due to the use of organochlorine pesticides, but that the effective population size was low and genetic monitoring is needed (Ponnikas et al. 2017). Nesje et al. (2000), however, based on 12 microsatellite loci, found lower heterozygosity values in Scottish, Scandinavian, Canadian, American and Tasmanian populations of peregrine falcons than those reported here, but the Scottish population had similar allelic richness (and one monomorphic locus) to the UK populations described in this study, and all the other peregrine populations had higher allelic richness.

Island (*F. p. nesiotes*) and mainland Australian (*F. p. macropus*) peregrine populations in the South Pacific and mainland and island populations in the North Pacific (*F. peregrinus*) described in Talbot et al. (2011) were also less genetically diverse than the peregrine populations described here, but peregrine populations from Vanuatu (South Pacific) and Colville area of northern Alaska showed higher number of private alleles than the European peregrines. Similarly, Talbot et al. (2011) found much lower mtDNA haplotype and nucleotide diversities in the South Pacific islands than in our study but their results from the North Pacific islands were comparable. There was also significant genetic differentiation with both molecular markers among their sampled populations in the South Pacific and elsewhere, suggesting isolation and long-term residency due to strong philopatry by peregrines, which reduces the capacity for gene flow and a limited ability for long-distance dispersal (Talbot et al. 2017).

There are several possible intrinsic and extrinsic factors, such as dispersal (natal and breeding), life history, behaviour, habitat and connectivity (Kozakiewicz et al. 2017), influencing the genetic composition of the newly re-established Sussex population. It is possible that not enough time elapsed for genetic erosion to occur or to be observed after the population bottleneck (Hailer et al. 2006; Johnson et al. 2010; Ponnikas et al. 2017). Studies has shown that longevity is instrumental to reducing genetic drift working as an intrinsic buffer against loss of genetic diversity (Hailer et al. 2006; Ponnikas et al. 2017). Long generation time is characteristic of many raptor species, as seen in White-tailed eagles (Hailer et al. 2006) and Golden eagles (Sonsthagen et al. 2012), often extending over twenty to thirty years. Comparably, peregrines in the wild have also been known to reach up to sixteen years of age, although mortality is typically mostly at 6-8 years (Tordoff and Redig 1997; London Peregrine Partnership).

### The origin of the Sussex peregrine population

Our phylogenetic analyses based on mtDNA was unable to detect the ancestral origins of the re-colonised population of peregrines; however, microsatellite data detected signatures of genetic structure demonstrating that the contemporary wild peregrine population in Sussex is genetically similar, but not identical, to the pre-pesticide UK population, and that it is genetically different from other European populations and from the domestic stock.

Despite UK peregrines possessing no migratory behavioural traits, it is a species of innate dispersal and colonising capability (Smith et al. 2015; McGrady et al. 2017; Ponnikas et al. 2017). Individuals dispersing from other refugial areas, such as Cornwall, Cumbria, Wales and Scotland (Crick and Ratcliffe 1995; Horne and Fielding 2002; Franklin and Everitt 2009), but genetically similar to the historical population, could have contributed to the Sussex peregrine population – however, we lack the necessary samples to test this hypothesis. Mature female peregrines may have contributed to the new population as some have been observed to change territories each year, rather than returning year on year to the same breeding territories (Mearns and Newton 1984). It is possible that non-breeders (floaters), dispersing but sexually inactive birds and migrants from regions not severely affected by organochlorine pesticides had an effect on the structure, dynamics, persistence and recovery of peregrine falcon populations, as has been hypothesised for other avian populations (Penteriani et al. 2011), and might have been crucial for the persistence of peregrine falcons regionally in the UK, rather than locally in the cliff areas of Sussex. Further sampling of wild peregrine populations from northern Europe and from the Irish and British Isles is needed, especially Wales and Scotland, as well as a more comprehensive sampling of domestic peregrines and wild subspecies of *F. peregrinus*.

Based on microsatellite data, we reason that the domestic falcons have not contributed to the genetic diversity of UK peregrines or to the re-establishment of the Sussex peregrine population. There was significant genetic differentiation between the Domestic group and the historical and contemporary wild UK peregrines, the Bayesian clustering method showed that the Domestic group was genetically different from the UK and other European peregrine groups, and the genetic assignment test did not place any UK-post peregrine in the Domestic group. The analysis of mtDNA data was less conclusive than with microsatellites due to the restricted number of nucleotides and low haplotypic diversity, as shown in other studies on peregrine falcons (e.g. Brown et al. 2007; Talbot et al. 2011; White et al. 2013). The only two haplotypes found in the Domestic group were also found in all other peregrine groups, and the UK-post population had two haplotypes not found in the Domestic group. If mtDNA from the domestic falcons (from other species of falcons or subspecies of *F. peregrinus*) had introgressed the contemporary population of wild UK peregrines, we would have expected to find more haplotypes in UK-post, or at least more shared haplotypes between Domestic and UK-post. MtDNA might not be the best marker to explore introgression due to the low mtDNA genetic variation in falcons in general (e.g. Brown et al. 2007; Talbot et al. 2011; White et al. 2013), as expected from recently diversified falcons (Nittinger et al. 2005; White et al. 2013).

Other studies using microsatellite markers have suggested introgression of alleles due to re-introductions of individuals from distant geographic origins (Tordoff and Redig 2001; Brown et al. 2007), but in this study there was a clear genetic distinction of domestic, Mediterranean falcons (*F. p. brookei*) and gyrfalcons from the wild peregrine falcons; however, further studies looking at introgression from domestic falcons and peregrine falcon subspecies into the recovering population of wild peregrine falcons (*F. p. peregrinus*) using a more comprehensive sampling scheme and possibly reduced representation DNA libraries (e.g. via Restriction site Associated DNA sequencing – RAD-seq) are needed.

### Implications for the protection of peregrines in the UK

Currently the global populations of peregrines have made a successful recovery and the population trends appears to be stable (BirdLife International 2016), and in the UK the peregrine falcon has subsequently been moved from amber to green on the Birds of Conservation Concern (Eaton et al. 2015). In Sussex, the newly established peregrine falcons are currently occupying all known historical nesting territories within the county, and other inland non-traditional sites, indicating retention of good-quality habitat, with no significant barriers to dispersal.

We recognise that there are implications for the protection of UK peregrines if it is accepted that there has not been a genetic bottleneck effect, and that contemporary and pre-pesticide populations contain similar levels of genetic diversity, as described here. Reestablishment of the population has been rapid, with conservation of the integrity of its genetic architecture without reintroductions from captive breeding and release schemes such as has occurred in North America, Scandinavia and Germany. We would suggest that for Sussex peregrines, and indeed other adjacent UK populations, there is no need for genetic conservation by wild-take and subsequent captive breeding programmes, and that the various legal conventions for peregrines already in place (e.g. UK Wildlife and Countryside Act 1981) and protection of breeding sites should be enough to prevent a new population decline in this recently recovered avian population. Although it appears that population growth rate of long-lived species (like the peregrine falcon) is more sensitive to changes in survival than to reproductive success (Smith et al. 2015; McGrady et al. 2017), any harvest of different age classes, if allowed by law, should be scientifically informed to assert minimal impacts to wild populations, and that wild populations should be adequately monitored to determine any potential impact in population demography and survival (Millsap and Allen 2006).

Furthermore, it appears that the loss of domestic birds (presumably of mixed origins) has not influenced the genetic structure of UK peregrine falcons. Domestic peregrines have been found to be genetically different from wild peregrines (at least as supported by the Bayesian clustering analysis), therefore not supporting the hypothesis that domestic birds could have contributed to the reestablishment of the Sussex peregrine population. Nonetheless, continuous loss of domestic birds could eventually lead to the introduction of alleles and mtDNA haplotypes not typically belonging to *F. p. peregrinus*, and careful monitoring and care while carrying out falconry should be advocated.

Although the estimated effective population sizes of peregrines studied here were found to be below the critical threshold of 50 individuals to avoid short-term inbreeding depression (Brook et al. 2002), at least in Sussex there is a healthy breeding population that could safeguard its long-term survival. However, it is recommended that field monitoring is carried out on a yearly basis to assess the Sussex peregrine population, and that genetic monitoring should be performed at 8 to 10-year intervals, in-line with generation lifespans of this species in the wild, to ensure that the evolutionary potential is not being compromised by inbreeding depression and genetic drift through intrinsic and anthropogenic factors.

## Acknowledgements

Several tissue samples were kindly provided by or obtained from museums, including Booth Museum Brighton, Natural History Museum at Tring, The World Museum (Liverpool) and Leeds Museums and Galleries. Many feather samples were collected by members of the public, and we acknowledge the help provided by bird breeders. German samples were kindly provided by Dr Peter Wegner from his North Rhine-Westphalia study population. Throughout the years, feather samples of Sussex peregrines were collected by The Sussex Peregrine Study. Tanya Thompson performed initial mtDNA sequencing of several peregrines used in this study as part of her undergraduate project at CCCU. Internal funding was provided to RV (CCCU) and to HJ and AW (UKC), and external funding was provided by The Sussex Peregrine Fund.

## Electronic supplementary material

### ONLINE RESOURCE 1

#### Comparative population samples

##### German

The German peregrine population was also extirpated during the 1950’s and 60’s with only around 40 pairs remaining, these in the south of the country (Saar 1988). The recovery in North Rhine–Westphalia and other German regions was assisted by the reintroduction of just over 1000 captive bred birds comprising some *F. p. brookei* but predominantly *F. p. peregrinus* (Saar 1988; Wink 2019) and it is currently considered a hybrid zone of between the two (Michael Wink, pers. comm.). Wegner et al. (2009) speculates that the North Rhine-Westphalia population underwent a genetic bottleneck and predicted low genetic diversity, confirmed using microsatellites (Volkhausen 2007).

##### Ireland

As with mainland UK the peregrine population of the island of Ireland (i.e. Republic of Ireland and Northern Ireland) suffered a population decline in the 1960’s followed by a rapid re-establishment (Ratcliffe 1984; Norriss 1995; Madden et al. 2009). We could find no records that any captive breeding and release programme was involved in the recovery of the Irish population. Moreover, among western European countries the Republic of Ireland is unique in that it allows wild take of peregrines for falconry purposes (Ryan 2016). Any falconry losses therefore would be of indigenous stock, and because there are few falconers in Ireland as compared to the UK, the number of losses would be minimal.

##### Domestic

We regarded domestic peregrines as those falcons available for falconry and advertised as ‘peregrines’ on the open market in the UK, thus freely available to falconers and as such may potentially be lost and breed in the wild. Registration of peregrines with the UK Department for Environment, Food and Rural Affairs (DEFRA, formerly the UK Department of Environment, DoE) by falconers and breeders was compulsory during the period 1985-2007 and this seems to be the period when captive breeding became prolific. Some registration data are available for this period (Fox and Chick 2007; Flemming et al. 2011) and from these inferences on the origins and size of the UK Domestic population can be made. During the period 1985-2007 some 627 peregrines of wild UK origin (wild-disabled and licensed wild-take) were held in captivity and of these, 130 (21%) produced offspring. Additionally, 398 peregrines were imported into the UK during the period 1976-2005 and were of subspecies other than the native *F. p. peregrinus*, namely *F. p. anatum, F. p. brookei, F. p. calidus, F. p. pealei* and *F. p. peregrinator*. If we assume that these imports had a similar breeding success rate as captive wild UK peregrines, we conclude therefore the founder breeding population for domestic birds was around 200-250 breeding birds and of these approximately 2/3 were native *F. p. peregrinus* and 1/3 imported birds of other subspecies.

According to Flemming et al. (2011) for the 22-year period of compulsory registration (1985-2007) 687 peregrines were lost by falconers, or roughly 30 per year. There is now no legal requirement to report lost birds but a voluntary registration scheme (The Independent Bird Register) provides a lost-found service to subscribers and some data indicates an almost identical estimate of 30 lost and not recovered peregrines per year (Barbara Royle pers. comm.). These two sources combined also provide evidence that similar losses of hybrids between peregrines and other falcon species. Therefore, in the 37 years since records began in 1983, around 1000 peregrines and a similar number of peregrine hybrids could have been lost into the wild by falconers. This figure is comparable to the number of captive bred birds purposely released in the German reintroduction scheme (Saar 1988) and indeed captive breeding/release schemes for other raptor species.

##### Falco peregrinus brookei

The Mediterranean peregrine (*F. p. brookei*) as its name suggests inhabits those regions of southern Europe on the fringe of the Mediterranean Sea (White et al. 2013). Its relative proximity to *F. p. peregrinus* populations in UK and mainland Europe makes it a potential candidate as a source of immigrants into depleted post-pesticide *F. p. peregrinus* populations. Further, Wink (2019) using mtDNA cytochrome b data suggests that western Europe represents a zone of integration between these two sub-species and intermediate populations have also been recorded along the French-Spanish Pyrenees (Zuberogoitia et al. 2009).

---

**Fig S1.**
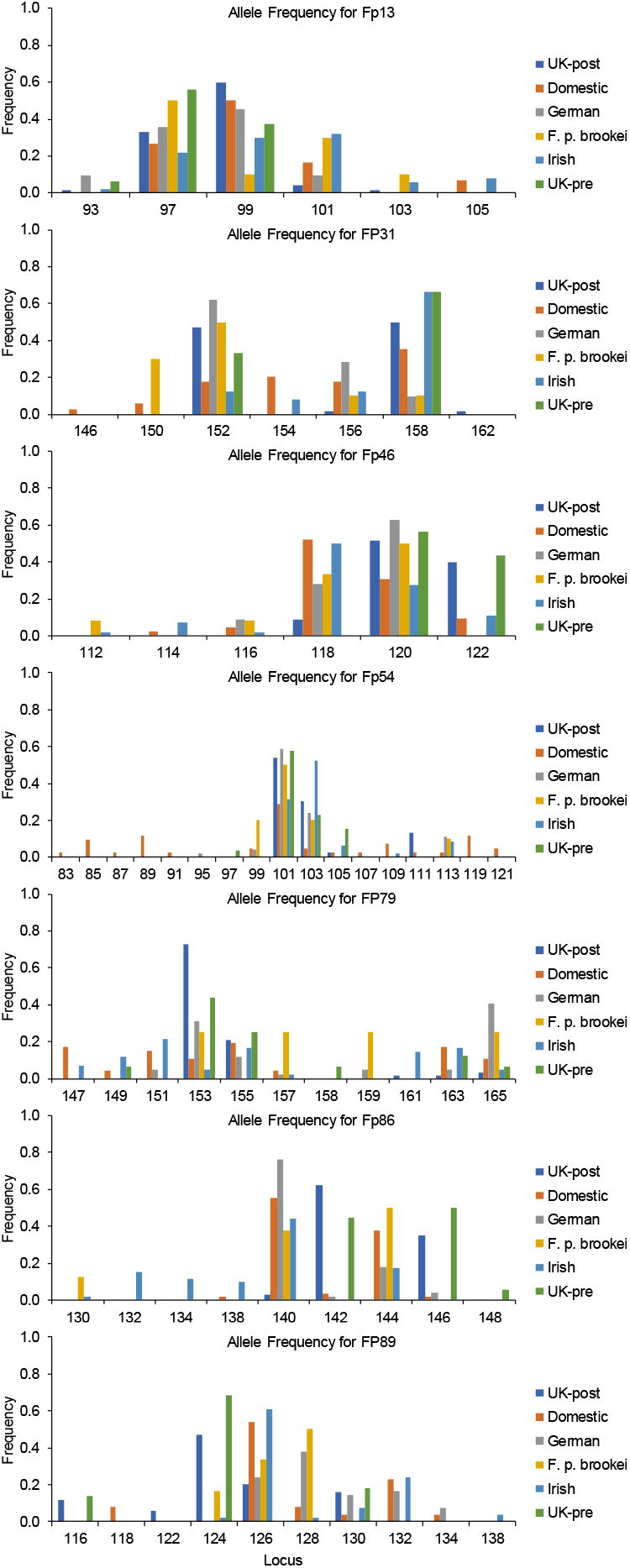

**Table S1.**
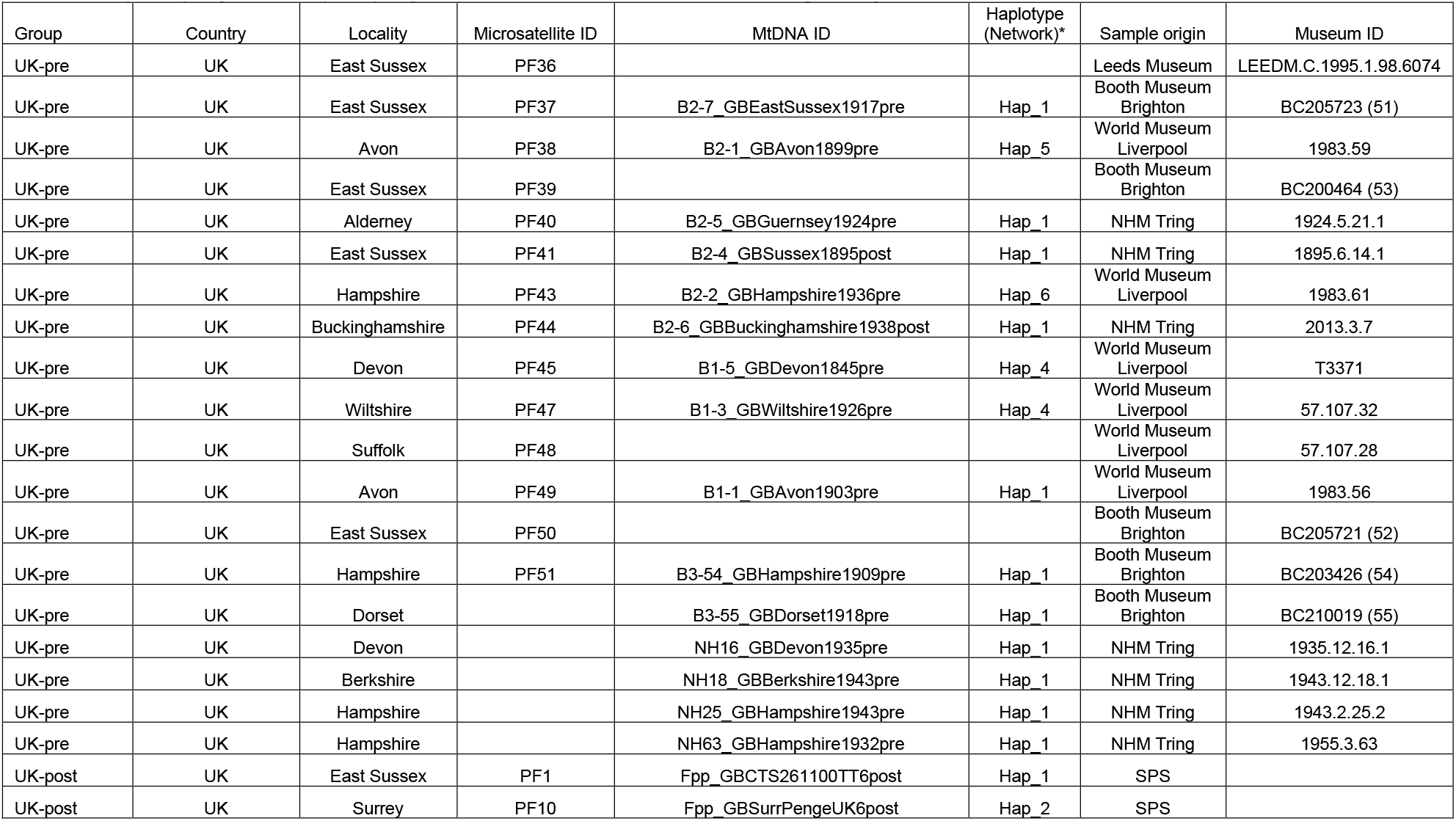
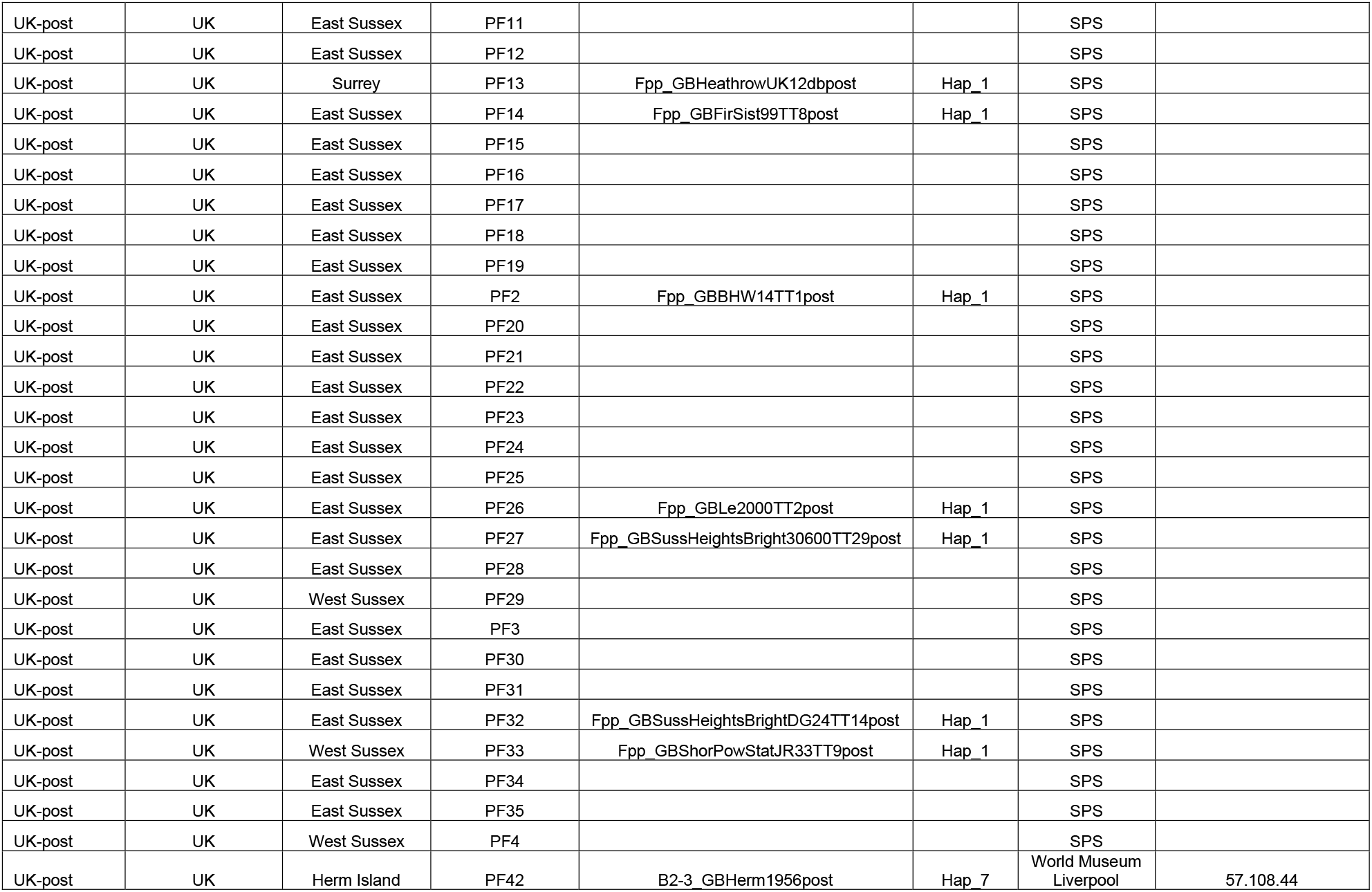
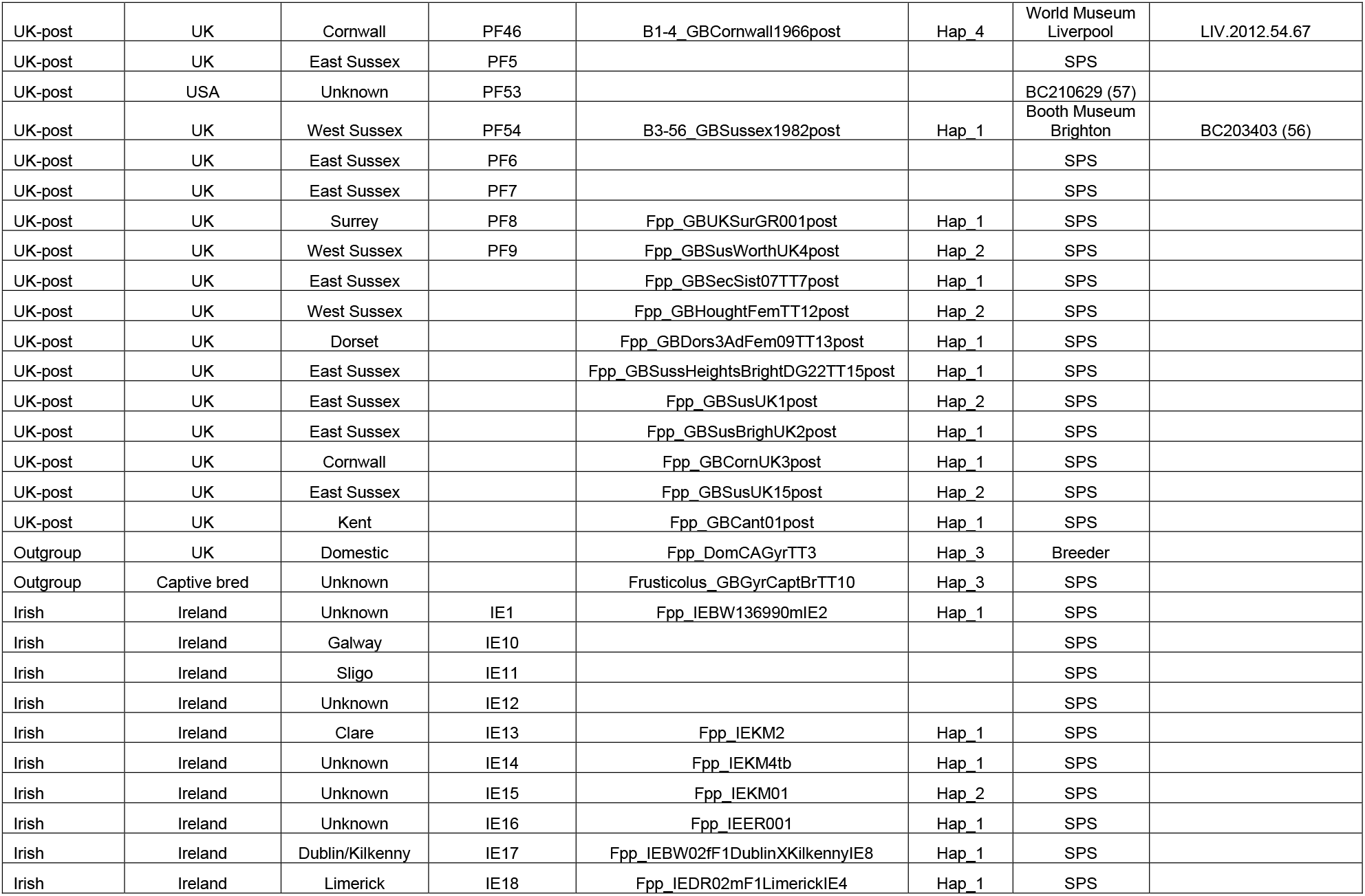
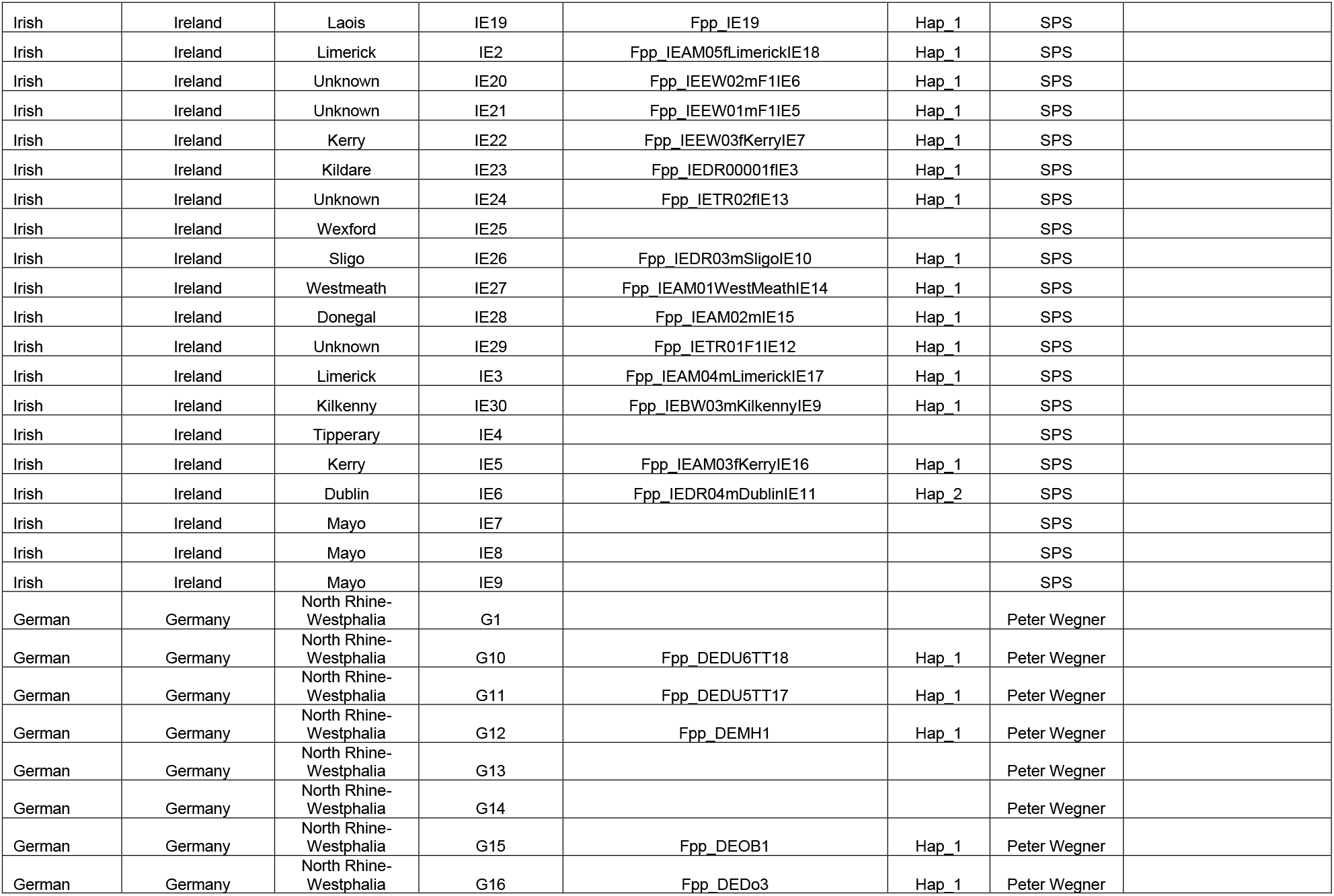
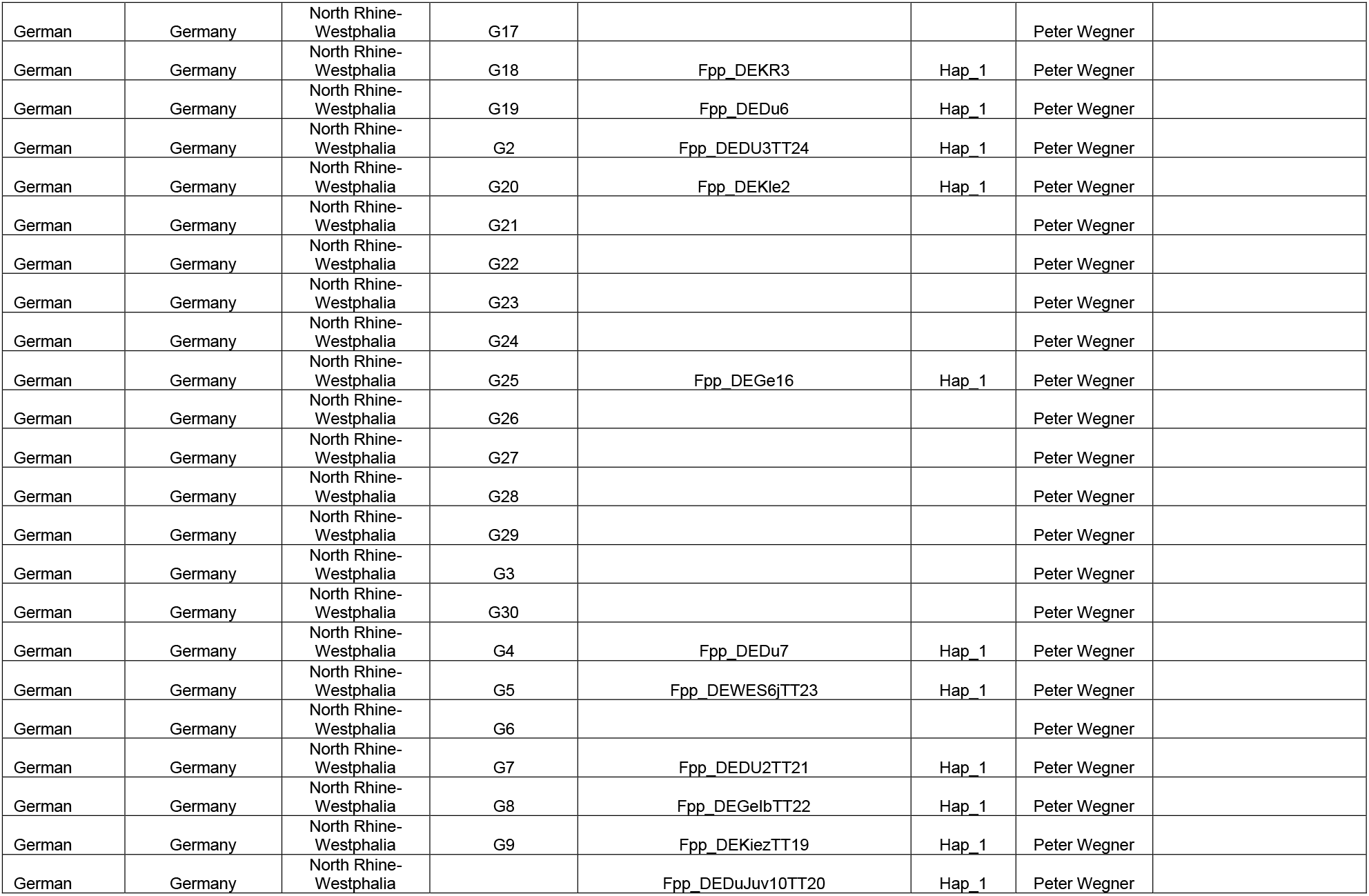
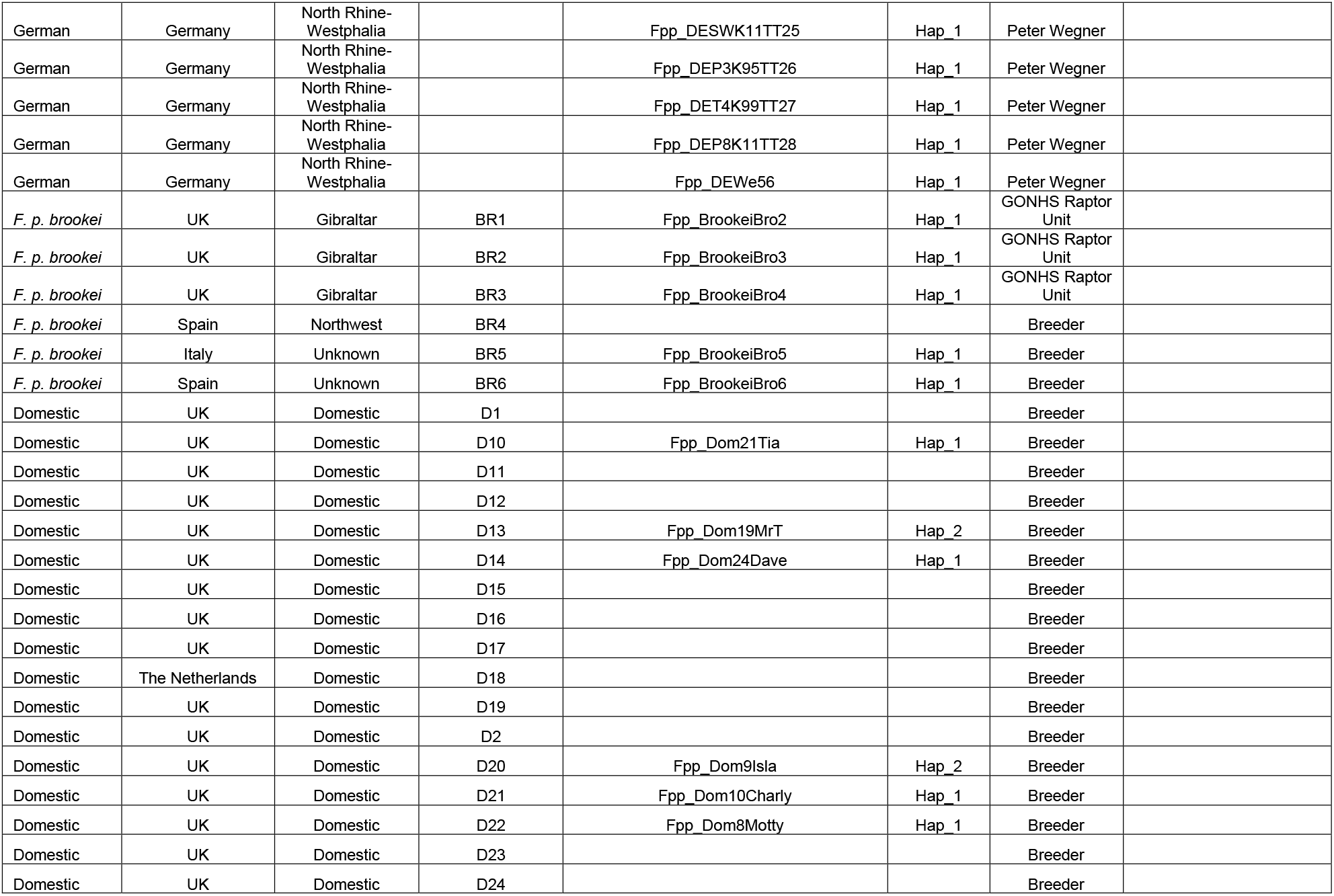
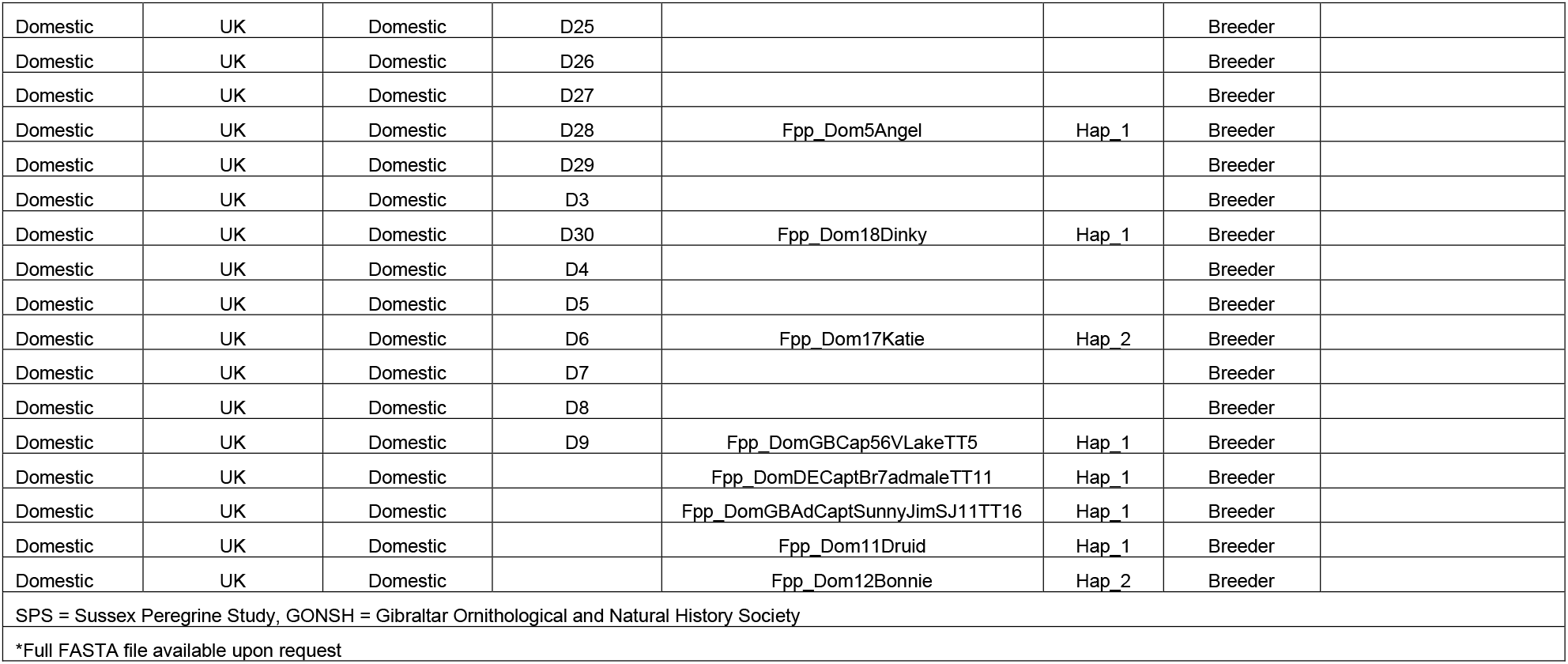
Samples of peregrine falcons (*Falco peregrinus*) used of microsatellite and mtDNA control region analysis

**Table S2.**
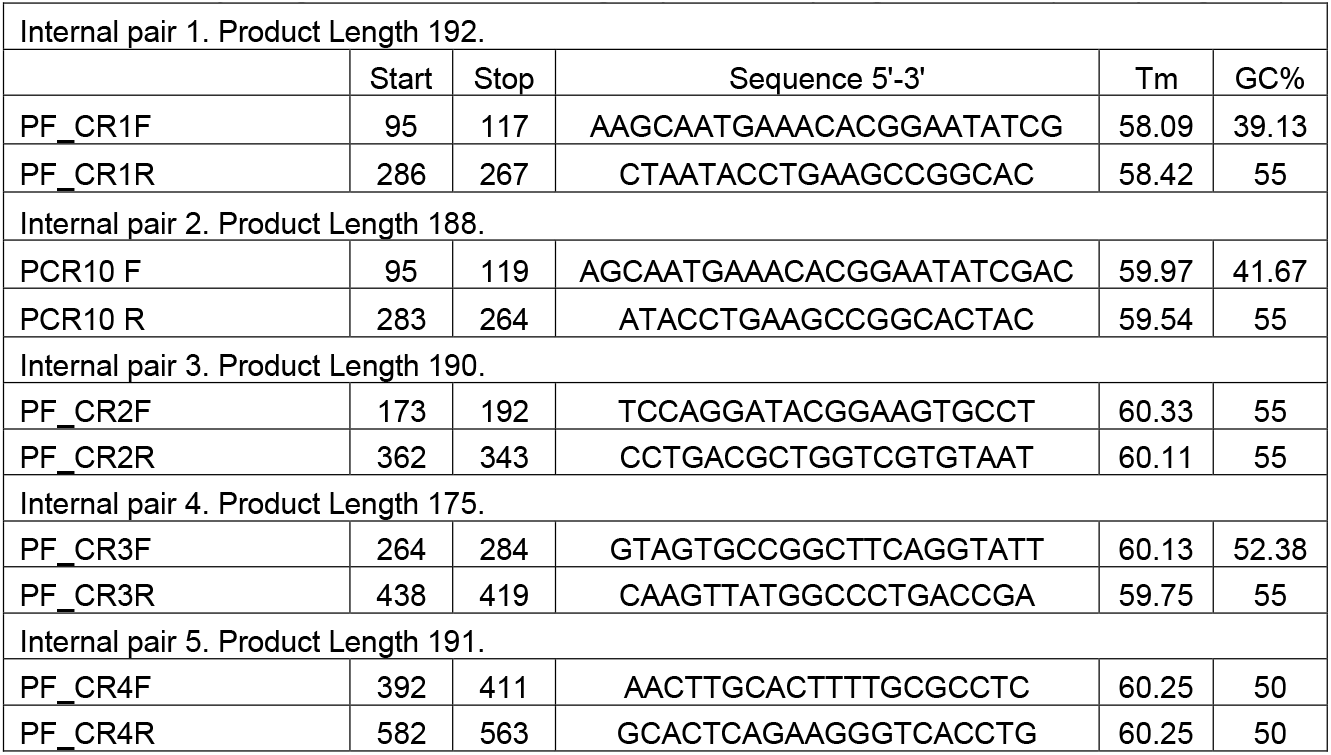
Newly designed mtDNA control region primers for peregrine falcons (*Falco peregrinus*)

**Table S3.** Nei’s unbiased pairwise genetic distances (above diagonal) and pairwise genetic differentiation (F_ST_) values (below diagonal) among peregrine falcon (*Falco peregrinus*) populations based on microsatellite data

|  |  |  |  |  |  |  |
| --- | --- | --- | --- | --- | --- | --- |
|  | UK-post | Domestic | German | *F. p. brookei* | Irish | UK-pre |
| UK-post | - | 0.591 | 0.459 | 0.469 | 0.591 | 0.020 |
| Domestic | **0.208** | - | 0.211 | 0.176 | 0.099 | 0.690 |
| German | **0.158** | **0.050** | - | 0.012 | 0.375 | 0.544 |
| *F. p. brookei* | **0.171** | **0.099** | 0.029 | - | 0.315 | 0.490 |
| Irish | **0.187** | **0.062** | **0.081** | **0.070** | - | 0.622 |
| UK-pre | **0.120** | **0.135** | **0.130** | **0.136** | **0.135** | - |
| Significant $F_{ST}$ values in bold ($P < 0.05$) | | | | | | |

**Table S4.** Pairwise genetic differentiation (F_ST_) values among peregrine falcon (*Falco peregrinus*) populations based on mtDNA control region data

|  |  |  |  |  |  |  |  |  |  |
| --- | --- | --- | --- | --- | --- | --- | --- | --- | --- |
|  | UK-post | UK-pre | German | Irish | GenBank-peregrines | GenBank-others | Domestic | *F. p. brookei* | *F. rusticolus* |
| UK-post | - |  |  |  |  |  |  |  |  |
| UK-pre | 0.050 | - |  |  |  |  |  |  |  |
| German | 0.124 | 0.026 | - |  |  |  |  |  |  |
| Irish | 0.008 | 0.029 | 0.048 | - |  |  |  |  |  |
| GenBank-peregrines | 0.089 | 0.050 | 0.107 | 0.065 | - |  |  |  |  |
| GenBank-others | 0.019 | 0.021 | 0.012 | 0.000 | 0.017 | - |  |  |  |
| Domestic | 0.000 | 0.108 | 0.231 | 0.056 | 0.145 | 0.060 | - |  |  |
| *F. p. brookei* | 0.124 | 0.026 | 0.000 | 0.048 | 0.107 | 0.012 | 0.231 | - |  |
| *F. rusticolus* | 0.838 | 0.814 | 0.931 | 0.899 | 0.880 | 0.859 | 0.843 | 0.931 | - |

